# Segmentation gene expression and function in *Vanessa cardui,* an emerging model for Lepidoptera

**DOI:** 10.64898/2026.06.01.729330

**Authors:** Ximena Gutiérrez Ramos, Katie Reding, Leslie Pick

## Abstract

Although all insects are segmented, the genes that control this process vary across species. Many of the pair-rule (PR) genes that direct segment formation in *Drosophila* are similarly utilized in other holometabolous insects, but more distantly related species use different genes for PR-patterning. Previously, we showed that Lepidoptera lack a highly conserved PR-gene, *paired.* Here, we used the painted lady butterfly *Vanessa cardui* as a lepidopteran model to explore the expression and function of PR-genes in this large clade of moths and butterflies. Orthologs of four *Drosophila* PR-genes are expressed in PR-like stripes and at least one displays PR-like function. Neither of the two genes that have PR-function in Hemiptera but not in *Drosophila* have PR-roles in *Vanessa.* Rather, the hemipteran PR-gene *Blimp1* functions in a novel fashion in abdominal segmentation in *Vanessa*. Thus, while butterflies appear to share PR-patterning mechanisms with other insects, they utilize only a subset of the *Drosophila* PR-gene orthologs and have not taken on hemipteran PR-orthologs for this process. These findings suggest extensive rewiring of the segmentation gene regulatory network in Lepidoptera.

## Introduction

The organization of the basic body plan into repeated segments along the antero-posterior (AP)-axis is shared across bilaterian animals (reviewed in Blair 2008; Diaz-Cuadros et al. 2021). This process has been studied in diverse taxa but, arguably, is best understood in the insect model species, *Drosophila melanogaster* (Wieschaus and Nüsslein-Volhard 2016). Sets of regulatory genes act in a well-studied hierarchy to generate and then diversify *Drosophila* body segments (Barresi and Gilbert 2023). Mutations in the pair-rule (PR) genes result in loss of alternate segment-wide units across the anterior-posterior body axis, indicative of a double-segment-wide prepattern early in development (Nüsslein-Volhard and Wieschaus 1980). In *Drosophila,* most PR-genes are expressed in 7-stripes in early embryos, in the primordia of the alternate segmental regions they specify (Barresi and Gilbert 2023). However, broadly expressed genes can still function as PR-genes; for example, *fushi tarazu factor 1* (*ftz-f1*) is expressed ubiquitously but its PR-function is restricted by the striped expression of its obligate partner Fushi Tarazu (Ftz) (Yu et al. 1997; Guichet et al. 1997; Yussa et al. 2001).

Although this segmentation gene network was worked out in *Drosophila*, it is reasonable to expect that this process is regulated by the same genes in other insects, given that all insects are segmented. However, there is significant variation in the timing of segment establishment across insects, which may suggest corresponding variability in genetic regulation. Segments in *Drosophila* are established more-or-less simultaneously at the blastoderm stage (simultaneous-segmentation). Many other insect species establish only the anteriormost segments in early blastoderm and add remaining segments one at a time as the germband elongates (sequential segmentation; reviewed in Davis and Patel 2002; Liu and Kaufman 2005). Is segmentation in sequentially segmenting insects regulated by the same gene network as in simultaneous segmenting insects?

This question has been most thoroughly addressed in beetles. In *Tribolium*, most PR-orthologs are expressed in striped patterns (Sommer and Tautz 1993; Brown et al. 1994; Patel et al. 1994; Brown and Denell 1996; Maderspacher et al. 1998; Choe et al. 2006; Choe and Brown 2007; Aranda et al. 2008; Janssen et al. 2010; Heffer et al. 2013; Choe et al. 2017; Clark and Peel 2018). RNA interference (RNAi) knockdown of *Tribolium even-skipped (eve)*, *runt* (*run*) or *odd-skipped* (*odd*) resulted in truncated embryos that displayed little more than a head, suggesting that these genes are required for germband elongation (Choe et al. 2006). Similar head-only phenotypes were obtained in a second beetle species, *Dermestes maculatus* (Xiang et al. 2017). However, weaker RNAi knockdown revealed segmental fusions and deletions in PR-registers for these genes (Xiang et al. 2017). Thus, in beetles, strong knockdown of three PR-genes resulted in inhibition of germband elongation and produced a truncated embryo, while weaker knockdowns that allowed germband elongation to proceed revealed *Drosophila*-like PR-defects. In *Tribolium*, only two PR-mutants were isolated in a mutant screen, corresponding to *paired* (*prd*) and *sloppy paired* (*slp*), the two genes with exclusively PR-like defects after RNAi knockdown in *Dermestes* (Maderspacher et al. 1998; Choe and Brown 2007; Xiang et al. 2015; Xiang et al. 2017). Variation in expression and/or function in either one or both beetle species was found for other PR-orthologs (Stuart et al. 1991; Aranda et al. 2008; Heffer et al. 2013; Choe et al. 2017; Xiang et al. 2017; Clark and Peel 2018). Collectively, these results suggest that while much of the regulatory network coordinating segmentation in *Drosophila* is conserved in sequentially segmenting beetles, it is not precisely recapitulated.

More limited studies have been carried out in other holometabolous insects, including the wasp *Nasonia vitrippenis* and the honeybee, *Apis mellifera,* where several PR-orthologs have taken on additional roles (Binner and Sander 1997; Osborne and Dearden 2005; Dearden et al. 2006; Keller et al. 2010, Wilson et al. 2010, Wilson and Dearden 2012; Rosenberg et al. 2014; Taylor and Dearden 2022 and see Discussion). In the mosquito *Anopheles stephensi*, the *prd* gene has been lost, but the *Pax3/7* family member *gooseberry* (*gsb)* has taken on PR-like expression and functionally replaced *prd* (Cheatle Jarvela et al. 2020), a clear example of how gene loss can be permitted after functional redundancy is established. In previous phylogenetic analysis of the *Pax* genes, we showed that *prd* is also absent from lepidopteran genomes (Cheatle Jarvela et al. 2020; Gutiérrez Ramos and Pick 2025). However, *gsb* was found in Lepidoptera. In fact, *gsb* appears to be conserved across all arthropod species, thus more broadly conserved than *prd* (Gutiérrez Ramos and Pick 2025).

Less is known about segmentation in the order Lepidoptera, butterflies and moths. Lepidopterans appear to form most of their segments sequentially from a posterior segment addition zone (SAZ) (Kraft and Jackle 1994, and see below). Some *Drosophila* PR-gene orthologs have been studied in the moths *Bombyx mori* and *Manduca sexta*, revealing conservation of PR-expression and function for some PR-orthologs (see Discussion) but we have not found reports of PR-gene expression in early embryos of any butterfly.

Here, we used techniques that we previously established in the painted lady butterfly, *Vanessa cardui* (*Vanessa, Vcar*) (Gutiérrez Ramos and Pick 2025), to examine PR-gene expression and function in lepidopterans. We found that *Vanessa gsb* is expressed and functions in stripes in all segments, similar to *Drosophila* (Nüsslein-Volhard and Wieschaus 1980; Bopp et al. 1986; Baumgartner et al. 1987). Orthologs of four PR-genes have PR-like expression in *Vanessa* and RNAi knockdown of *Vcar*-*eve* or *Vcar*-*run* revealed PR-like function for these genes. We also examined *Vanessa E75* and *Blimp1,* genes that function as PR-genes in the hemimetabolous insect *Oncopeltus fasciatus* (Erezyilmaz et al. 2009; Reding et al. 2024). RNAi knockdown of *Vcar-Blimp1* revealed a novel phenotype, with deletion of two abdominal segments. Overall, *Vanessa* appears to utilize only a subset of the PR-gene orthologs found in other insects for PR-patterning. How *Vanessa*, and possibly other Lepidoptera, pattern segments with a more limited or divergent set of genes remains to be determined.

## Results

### *Vanessa cardui* early embryo development

We examined the timing and stages of *Vanessa* development from egg laying to 16 hours after egg laying (AEL). *Vanessa* eggs are fertilized as they are laid (Figure 1Ai). Nuclear divisions start in the most anterior part of the egg (arrowhead, Figure 1Ai) and later distribute across the embryo. The first nuclear divisions were observed between 0-4 hours AEL (Figure 1Aii, 1Ci) and continued through 6 hours AEL (Figure 1Aiii-iv). The blastoderm formed at ∼8 hours AEL (Figure 1Av, 1Bi), characterized by a broad sheet of cells along the egg’s circumference (Figure 1Bi, 1Cii). Extra-embryonic tissue is distinguishable by large nuclei in anterior and posterior poles (Figure 1Bi, arrowheads). Next, the blastoderm compressed laterally, initiating germ layer formation within the germ primordium; the anterior margins of the blastoderm invaginated to form the head lobes. At this stage, the shape of the primordium resembled a saddle, with the segment addition zone (SAZ, S) at the posterior end (Figure 1Bii, 1Ciii, S). When the germband started to elongate, the protocephalon (Figure 1Biii-iv, 1Civ, asterisks) and the primitive midline groove were distinguishable. The germband continued to elongate, with segments added sequentially until ∼16 hours AEL, when the gnathal segments, the head lobes and the midline were evident (Figure 1Bv, 1Cv-vi).

**Figure 1.**
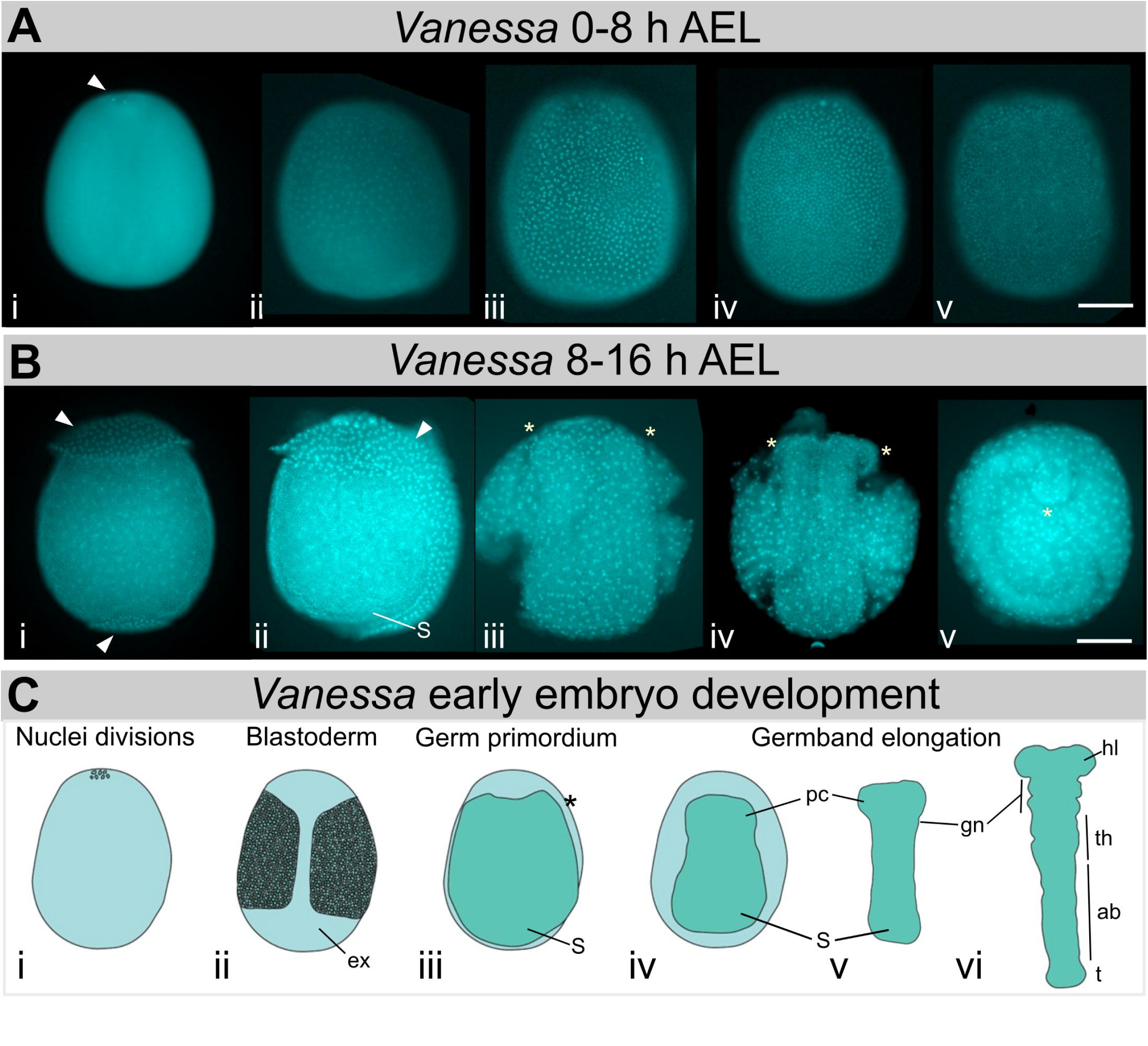
*Vanessa* early embryo development. (**A**, **B**) Hoechst nucleic acid staining of embryos at successive stages of development. (**A**) 0 to 8 hours AEL, (**B**) 8 to 16 hours AEL. (**C**) Schematic of the main developmental changes across the first 16 hours of *Vanessa* embryo. SAZ (S), protocephalon (pc), head lobes (hl), gnathal segments (gn), thoracic segments (th), abdominal segments (ab), telson (t). Images orientated anterior, top; posterior, bottom. Scale bar 200 um.

This early embryonic developmental progression is like that seen for other Lepidopterans such as *Bombyx* (Krause and Krause 1964; Miya 1985; Nagy et al. 1994), *Manduca* (Dorn et al. 1987; Broadie et al. 1991; Kraft and Jackle 1994) and *Bicyclus* (Holzem et al. 2019). However, we were unable to distinguish germinal disc formation that is localized in the anterior of *Bicyclus* embryos at 5 hours AEL (Holzem et al. 2019). Our experiments defined the timing of blastoderm formation (∼8 hours AEL), likely the earliest stage at which PR-gene expression would be expected.

### *Vanessa-gsb* is a segment polarity gene

We previously showed that *Vcar-gsb* is expressed in stripes with single-segment periodicity during germband elongation (Gutiérrez Ramos and Pick 2025). As *gsb* was found to have PR-function in a mosquito (Cheatle Jarvela et al. 2020), we asked if *Vcar-gsb* has also taken over PR-function from *prd*, or if it has segment polarity function, as in *Drosophila*, or some other function? We expanded our sampling of developmental stages to characterize *Vcar-gsb* expression from 8 to 16 hours AEL. The earliest pattern detected was two anterior stripes (Figure 2Ai, arrowheads). With the separation of the protocephalon (asterisks), the anteriormost stripe split medially and additional stripes were observed in the primordium of each segment (Figure 2Aii). No stripes were observed in the SAZ (Figure 2Aii-iii). As the germband elongated, each stripe appeared to split medially (Figure 2Aiv-v), as previously observed (Gutiérrez Ramos and Pick 2025). Thus, *Vcar-gsb* is expressed in the blastoderm and through germband elongation with single segment periodicity.

**Figure 2.**
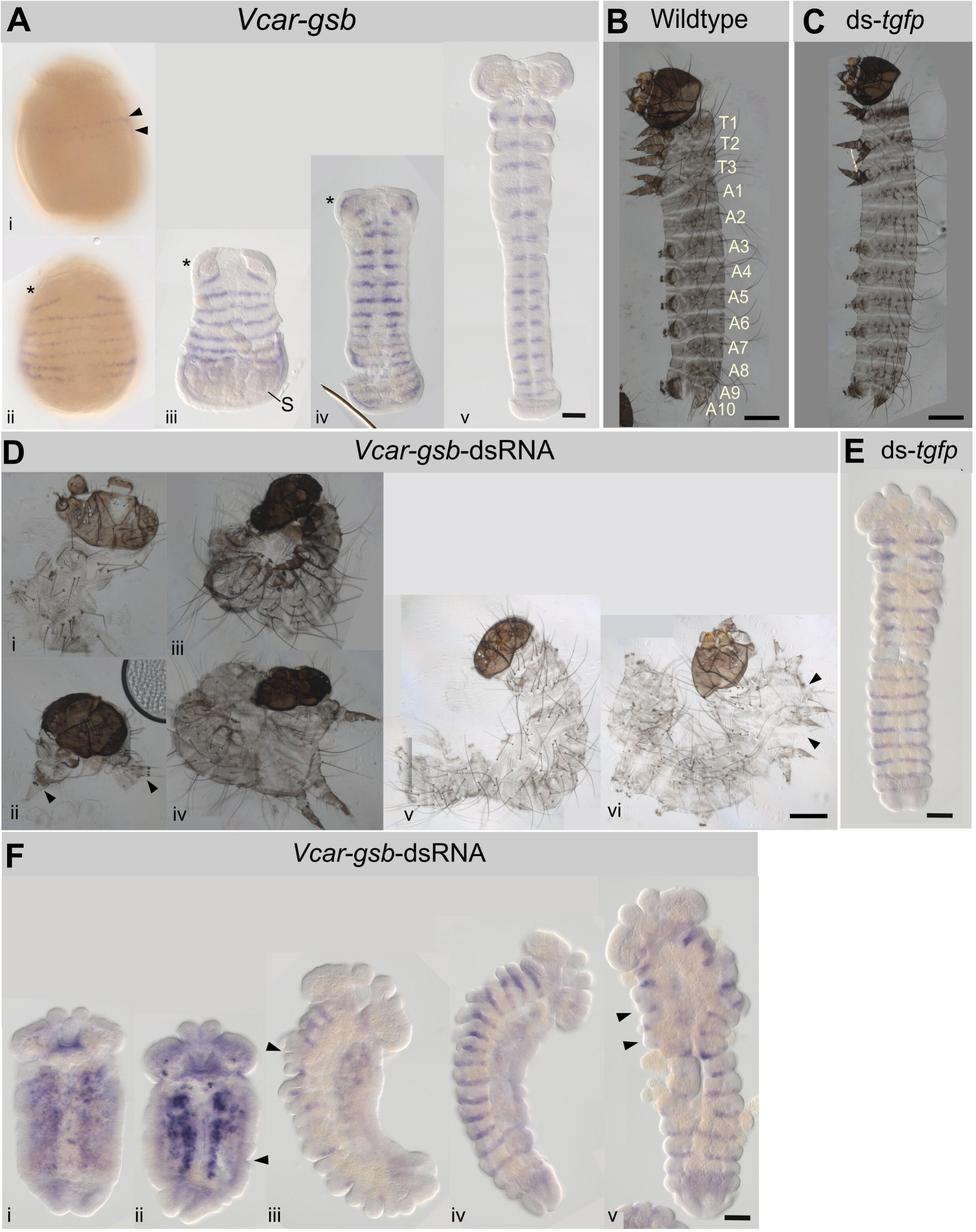
*Vcar-gsb* displays striped expression and segment polarity-like function. (**A**) Colorimetric *in situ* hybridization with *Vcar-gsb* probe at successive time points between 8 and 16 hours AEL. (i) Anterior blastoderm stripes (arrowheads). (ii) Multiple stripes in germ primordium, two most anterior stripes separating (asterisk). (iii) Early elongating germbands: segmental stripes; stripe in head lobes separated medially (asterisk); no expression observed in segment addition zone (S). (iv, v) Stripes in every segment with faint expression in head lobes (asterisk). (**B**) Cuticle preparation from *Vanessa* wildtype larva: head (top), three thoracic segments (T1-T3), ten abdominal segments (A1-A10). A3 to A6 and A10 each bear a pair of prolegs. (**C**) ds-*tgfp* eRNAi embryo displays wildtype-like phenotype. (**D**) *Vcar-gsb*-dsRNA eRNAi embryos: (i) head phenotype, (ii) head with one set of prolegs (arrowhead), (iii-iv) torus, (v-vi) partial fused segments. (**E-F**) *Vcar-en* in situ hybridization in (**E**) *tgfp*-dsRNA eRNAi embryo, (**F**) *Vcar-gsb*-dsRNA eRNAi embryos. Images orientated anterior, top; posterior, bottom. Scale bars: cuticle preparations, 200 um; in situ hybridization, 100 um.

We carried out embryonic RNAi (eRNAi) knockdown of *Vcar-gsb* using two non-overlapping double stranded RNAs (dsRNA), both targeting regions downstream of the homeobox (dsRNA1 and dsRNA2, Fig S1A). Embryos were injected with *Vcar-gsb*-dsRNA1, *Vcar-gsb*-dsRNA2, or *tgfp-*dsRNA (ds-*tgfp*) as a negative control, and allowed to develop until just before wildtype embryos would be expected to hatch (‘prehatchlings’). *Vanessa* wildtype prehatchlings have a head, three thoracic segments (T1-T3) and ten abdominal segments (A1-A10), with A3-A6 and A10 each possessing a pair of prolegs (Figure 2B, wildtype). After control ds-*tgfp* injection, the expected segments were observed in 86% of injected embryos (n=452/523, Fig. 2C). No development was observed for 13% (n=66/523) of injected embryos and 1% showed delayed development (n=5/523). Knockdown of *Vcar-gsb* resulted in four key phenotypes (Fig. 2D), comprising 85% n=384/450 of dsRNA1-injected embryos and 81% n=468/581 of dsRNA2-injected embryos (see details in Table S1): (1) The most severe phenotype, seen in ∼25% of prehatchlings, termed ‘head-like’ displayed the head capsule with stemmata, frons, and antennae similar to wildtype while mandibles, maxillae, and labrum presented appeared distinct from wildtype (Figure 2Di). (2) Less severe truncations, seen in ∼20% of prehatchlings, consisted of the head, undefined tissue, and one set of prolegs (Figure 2Dii, arrowheads). (3) ∼30% of prehatchlings displayed a torus-like shape, with thoracic and abdominal segments fused on just one side of the embryo, presumably causing the observed curvature (Figure 2Diii, ventral view; Fig. 2Div, dorsal view of different embryo). (4) Other prehatchlings had partial segmental fusions; in some cases A1-A4 were fused on one side of the embryo (Figure 2Dv); in others thoracic segments were fused on one side of the embryo (Figure 2Dvi), suggesting an intermediate form of the torus phenotype. Overall, segment formation defects were observed after *gsb* knockdown in more than 80% of embryos examined but no PR-like defects were observed, although we cannot rule out the possibility that the head-only phenotypes reflect a PR-like role.

Next, we asked if high concentrations of the dsRNAs masked potential early PR-function. We performed injections of *Vcar-gsb*-dsRNA1 at lower concentrations. The most severe phenotypes previously observed after knockdown with *Vcar-gsb*-dsRNA1 (head-like, head with one set of prolegs, and torus) were absent when 0.15 uM or 0.015 uM dsRNA was injected. Embryos injected with *Vcar-gsb*-dsRNA1 at 0.15 uM showed partial fused segments (41%, n=55/133), wildtype-like phenotypes (35%, n=47/133), developmental delay (19%, n=25/133), or no development (5%, n=6/133). At 0.015 uM, the frequency of partially fused segments decreased to 23% (n=33/142) and the frequency of wildtype-like increased to 64% (n=91/142) (Table S2). However, we did not observe a PR-like loss of alternate segments, although the specific identities of segments present in larvae with torus or with partial fusion phenotypes could not be definitively determined.

We also performed *in situ* hybridizations in *Vcar-gsb* dsRNA-injected embryos to visualize expression of *Vcar engrailed* (*Vcar-en*)—a conserved segment polarity gene (Wieschaus and Nüsslein-Volhard 2016). In ds-*tgfp* control embryos, *Vcar-en* stripes were present in every segment of elongated germbands (Figure 2E), as expected. In *Vcar-gsb*-dsRNA1 embryos, we observed a range of abnormalities, including embryos with head lobes and telson, but no defined segments (Figure 2Fi-ii) with *Vcar-en* expression concentrated into partially fused rows along each side of the midline, possibly due to fusion of multiple stripes along the AP-axis and/or failure of germband elongation. Some embryos displayed a well-defined head, telson and lateral segments, with some fused segments (Figure 2Fiii, arrowhead), with *Vcar-en* stripes visible in several segments. Other embryos completely lacked segmentation and *en* expression in one lateral half of the body, while displaying wildtype-like *en* stripes in every segment in the other half (Figure 2Fiv). Finally, some embryos displayed partially fused segments, with *Vcar-en* stripes in only some segments (Figure 2Fv). We did not observe a PR-like register to the segmental fusions in these embryos. Together, these findings suggest that *Vcar-gsb* is required for the formation of all the segments, similar to its role as a segment polarity gene in *Drosophila*.

### Some *Vanessa* pair-rule gene orthologs are expressed in pair-rule-like stripes

Given the lack of an obvious PR-role for *Vcar-gsb*, and the absence of *prd* from the genome, we asked if *Vanessa* use PR-patterning *per se* and if so, if orthologs of other *Drosophila* PR-genes drive this process. We identified *Vanessa* orthologs of *ftz*, *ftz-f1*, *eve*, *sloppy paired 1* (*slp1*), *sloppy paired 2* (*slp2*), *run*, *odd*, *hairy* (*h*) and *odd-paired* (*opa*) and performed *in situ* hybridization for each in 8-16 hours AEL embryos.

*Vcar-ftz* expression was observed as a wide band in the posterior part of the germ primordium (Figure 3Ai, arrowhead), followed by one anterior wide band, one posterior band, and expression in the SAZ (Figure 3Aii). Once the germband elongated, expression continued in the SAZ and midline of the germband (Figure 3Aiii). Later, expression appeared in the midline as spots in every segment, presumably in the developing nervous system (Figure 3A iv-v). *Vcar-ftz-f1* expression was not detected in the blastoderm or germ primordium (Figure 3Bi-ii) and later appeared to be faintly, ubiquitously expressed (Figure 3Biii-v).

**Figure 3.**
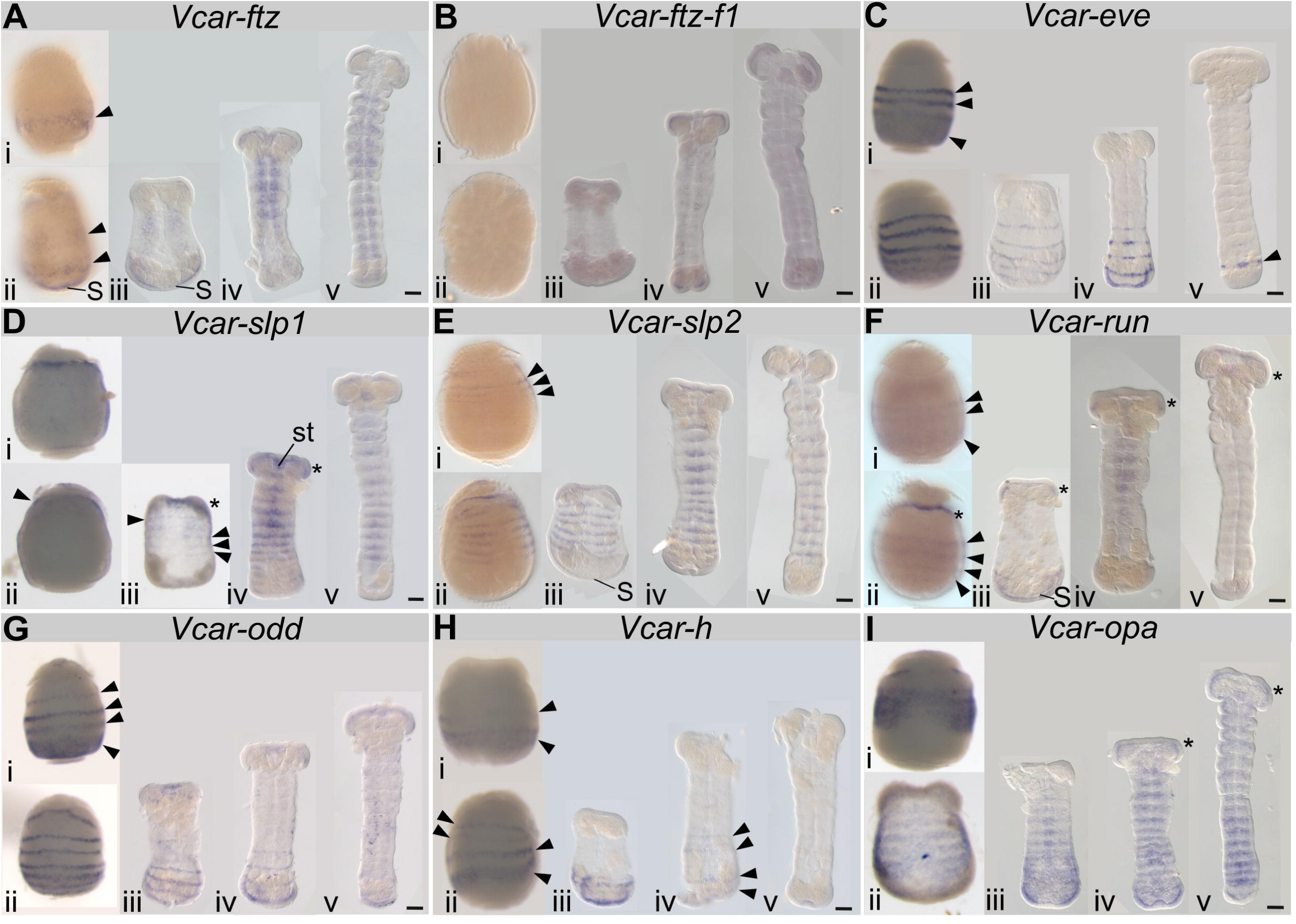
Expression patterns of *Vanessa* pair-rule gene orthologs. Colorimetric *in situ* hybridization with *Vcar-PR*-genes probes, as indicated at (i-ii) blastoderm stage, germ primordia, (iii-v) successive stages of germband development. (**A**) *Vcar-ftz*: (i) posterior band (arrowhead); (ii) 2 widely-spaced bands (arrowheads) and SAZ expression. (iii-iv) Expression in midline, likely the developing nervous system. (**B**) *Vcar-ftz-f1:* (i-ii) not detected, (iii-v) faint ubiquitous expression in elongated germbands. (**C**) *Vcar-eve:* (i) 2 widely spaced anterior stripes and broad posterior band (arrowheads), (ii) 5 stripes along the germband, (iii) 4 stripes in the posterior of the embryo, (iv) expression at posterior SAZ in fully elongated germbands, (v) one stripe anterior to the SAZ (arrowhead). (**D**) *Vcar-slp1:* not detected in (i) blastoderm or (ii) germ primordia. (iii) Stripes along the elongating germband (arrowheads) that (iv-v) later are clear in every segment, head lobes (asterisk) and stomodeum (st). (v) Striped expression maintained in the thoracic and abdominal segments. (**E**) *Vcar-slp2:* (i) anterior stripes (arrowheads); (ii) additional stripes present and (iii-iv) segmental stripes; no expression observed in SAZ. (v) Stripes centrally localized. (**F**) *Vcar-run*: (i) 2 widely spaced anterior stripes and a broad posterior stripe (arrowhead). (ii) Additional stripes in germ primordium (arrowheads). (iii) Expression in head lobes (asterisk) and posterior stripes, likely the SAZ (S); (iv-v) expression in head lobes (asterisk), midline paired dots in gnathal and thoracic segments, stripes in SAZ. (**G**) *Vcar-odd*: (i) 3 widely spaced anterior stripes and posterior broad band (arrowheads). (ii) Widely spaced stripes along germ primordium. (iii-iv) Striped expression continued in posterior germband which (v) faded at the end of the 16 hours AEL. (**H**) *Vcar-h:* (i-ii) 2-3 widely spaced stripes in blastoderm (arrowheads); (iii) stripes at posterior elongating germband and SAZ (arrowheads). (iv) Stripes still detectable (v) by 16 hours AEL no expression observed. (**I**) *Vcar-opa:* (i) broad anterior band, (ii) 4 stripes along the germ primordium. (iii) Segmental stripes, expression in SAZ and along midline, (iv) expression in head lobes (asterisk), segmental stripes and SAZ. (v) Segmental stripes in abdominal segments. Images orientated anterior, top; posterior, bottom. Scale bar, 100 um.

*Vcar-eve* expression was first detected in the blastoderm as 2 anterior stripes and a wide band in the posterior (Figure 3Ci, arrowheads). Later, this posterior band appeared to split into three separate stripes (Figure 3Cii). *Vcar-eve* stripes in the germband were widely spaced (Figure 3Ciii-iv), especially as compared to *gsb* expression in embryos of similar age (compare Figure 3Civ to Figure 2Aiv). As the germband elongated, stripes were seen only in the most posterior part of the germband (Figure 3Civ). In older germbands, *Vcar-eve* expression was observed in a single stripe close to the telson (Figure 3Cv, arrowhead).

*Vcar-slp1* expression was not detected in the blastoderm (Figure 3Di). In the germ primordium, anterior expression was observed in the protocephalon (Figure 3Dii, arrowhead), head lobes (asterisk), stomodeum (st) and in stripes along the germband (arrowheads) (Figure 3Diii-iv). Stripes were present as the germband elongated, with apparent segmental register (Figure 3Div-v). *Vcar-slp2* striped expression first appeared in the anterior part of the blastoderm (Figure 3Ei, arrowheads), and later stripes arose along the germ primordium with no expression detected in the SAZ (Figure 3Eii-iii). During germband elongation, *Vcar-slp2* was expressed in centrally localized stripes marking each segment (Figure 3Eiv-v).

*Vcar-run* expression was observed in the blastoderm as two anterior stripes and a wide posterior band (Figure 3Fi, arrowheads). In the germ primordium, widely spaced stripes of expression were observed (Figure 3Fii, arrowheads), similar to *Vcar-eve* in a similarly staged embryo (compare Figure 3Fii to Cii). During early germband elongation, expression was observed in the head lobes, in stripes in the posterior of the germband and in the SAZ (Figure 3Fiii). Expression continued in the head lobes, with segmental centrally located stripes in the gnathal and thoracic segments (Figure 3Fiv). Later, faint expression continued in the head lobes and stomodeum (Figure 3Fv, asterisk).

*Vcar-odd* expression was first observed in the blastoderm as three stripes and a wide posterior band (Figure 3Gi, arrowheads), similar to *Vcar-eve*. This wide band later resolved into several stripes (Figure 3Gii). When the germband started to elongate, striped expression was only observed in the posterior of the embryos (Figure 3Giii-iv). Expression was not detected in later germbands (Figure 3Gv).

*Vcar-h* blastoderm expression was first observed in stripes in the posterior half of the embryo (Figure 3Hi, arrowheads). Later, four widely spaced stripes of varying intensity were observed (Figure 3Hii). At the beginning of germband elongation, *Vcar-h* expression was observed in the SAZ (Figure 3Hiii). The stripes faded in the elongated germband (Figure 3Hiv, arrowheads) and were undetectable in older embryos (Figure 3Hv).

*Vcar-opa* expression appeared in the blastoderm as an anterior wide band that resolved into stripes along the germ primordium (Figure 3Ii-ii). In the elongated germband, striped expression was observed, in addition to expression along the midline, and posteriorly in the SAZ (Figure 3Iiii). Later, expression was seen in the head lobes, in segmental stripes, and in the SAZ (Figure 3Iiv). At the end of 16 hours AEL, the expression in the head lobes, gnathal, and thoracic segments appeared weaker, while segmental striped expression in the abdominal segments continued (Figure 3Iv).

Based on these expression patterns we can classify *Vanessa*-PR-genes in groups: 1) widely-spaced stripes suggestive of PR-like expression for *Vcar-eve*, *Vcar-run*, *Vcar-odd* and *Vcar-h*; 2) segmental stripes for *Vcar-slp2* and *Vcar-opa,* with segmental stripes appearing later for *Vcar-slp1*; and 3) *Vcar-ftz*, which was difficult to classify with widely spaced stripes in the blastoderm and presumptive central nervous system expression. *Vcar-ftz-f1* expression was very faint in embryos and we were not able to isolate it from early embryonic cDNA.

### *Vanessa eve* RNAi suggests classical pair-rule function

We next selected two genes that have PR-like expression, *Vcar-eve* and *Vcar-run*, and one with segmental expression, *Vcar-slp*, to assess function. For knockdown of *Vcar-eve*, we designed two non-overlapping dsRNAs, with dsRNA1 targeting most of the homeobox and dsRNA2 targeting the end of homeobox to the end of the transcript (Fig S1D). *Vanessa* embryos injected with 1.5 uM ds-*tgfp* as negative control showed wildtype-like phenotypes in 86% (n=452/523) of embryos (Figure 4A, Table S1). We observed three distinct segmentation phenotypes following *Vcar-eve*-dsRNA knockdown, in total comprising 55% (n=299/540) of dsRNA1-injected embryos: (1) the PR-like loss of the even-numbered abdominal segments (Figure 4B). Segments were identified by the presence of only one pair of legs (identifying T2) and the presence of only two pairs of prolegs, identifying segments A3 and A5. (2) Head-like, similar to those seen after knockdown of *Vcar-gsb* (Figure S2Ai). (3) Partial fusions, including loss of two thoracic segments, presumably T1 and T3, and one abdominal segment with three sets of prolegs remaining (Figure S2Aii, arrowheads). Others displayed fusion of thoracic segments, leaving just three legs (Supplementary Figure S2Aiii-iv, dots), and all abdominal segments (arrowheads). Partial fusions suggest an intermediate form of the PR-like phenotype. These phenotypes were seen for both dsRNAs, comprising 59% (n=267/450) of dsRNA2-injected embryos (Table S1). These results suggest that *Vcar-eve* has PR-function in *Vanessa*.

**Figure 4.**
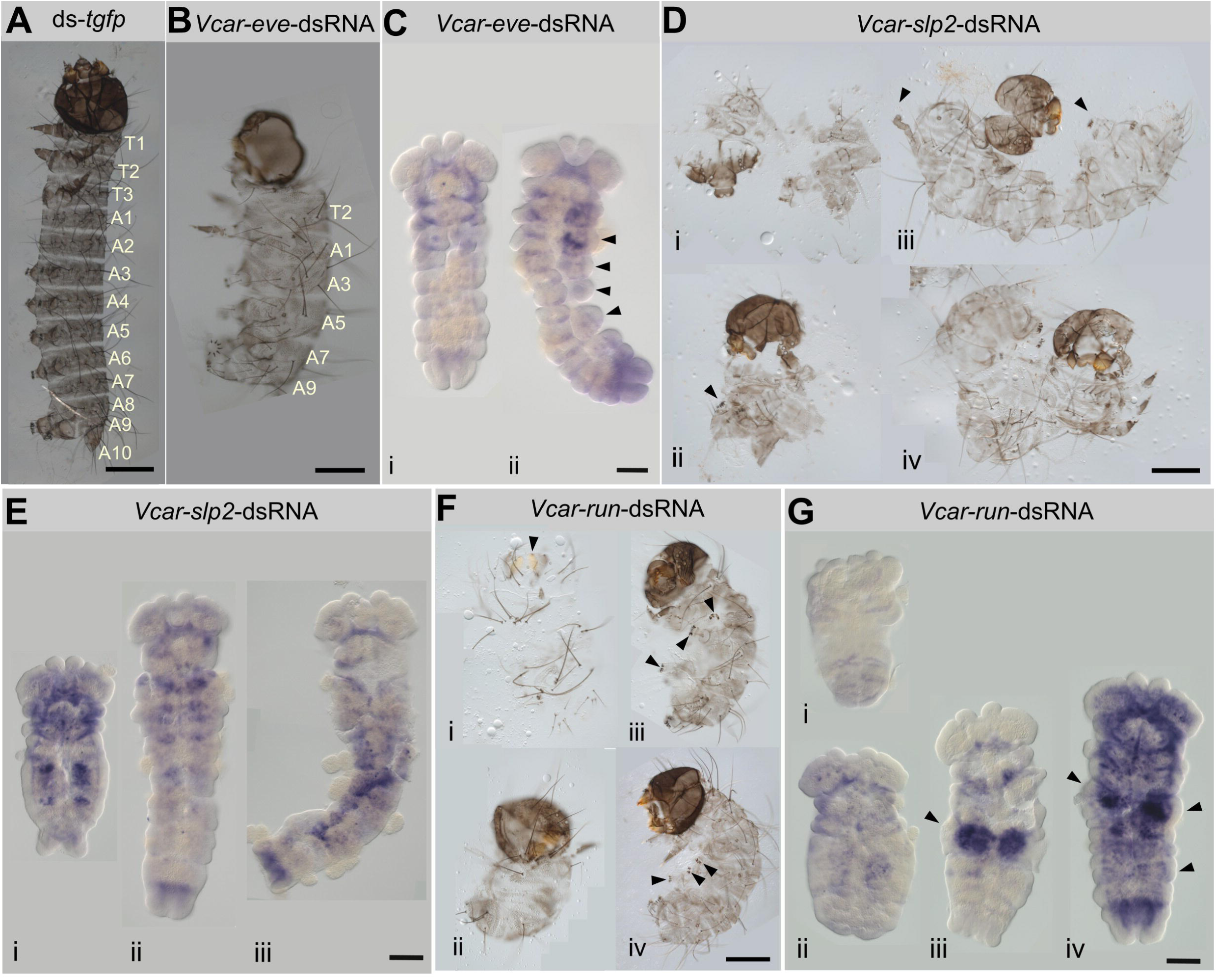
*Vanessa* pair-rule gene orthologs function in segmentation. eRNAi results: (**A**, **B**, **D** and **F**) cuticle preparations, (**C**, **E**, **G**) *Vcar-en* expression in eRNAi embryos 24 hours post-injection. (**A**) *tgfp*-dsRNA eRNAi embryos appeared wildtype-like. (**B**, **C**) *Vcar-eve*-eRNAi (**B**) alternate segments were lost; head, one pair of legs (T2) and five abdominal segments (A1, A3, A5, A7 and A9) were present. (**C**) (i-ii) Loss of segments and partial fusions (arrowheads) in germbands. (**D**, **E**) *Vcar-slp2*-eRNAi (**D**) (i) head-like; (ii) head and prolegs (arrowhead); (iii) head, legs (arrowhead) and prolegs (arrowhead); (iv) torus. (**E**) Examples of (i) embryos lacking defined segments and (ii-iii) developed germbands with non-specific *Vcar-en* expression. (**F**, **G**) *Vcar-run*-eRNAi (**F**) cuticle defects included (i) undefined mass; (ii) head and gnathal structures; (ii-iv) head, one leg and two or three prolegs (arrowheads). (**G**) (i) *Vcar-en* expression was faint and many germbands had undefined shapes; (ii-iii) defined head structures but undefined segments; (iv) fused segments (arrowheads). Images orientated anterior, top; posterior, bottom. Scale bars: cuticle preparations, 200 um; in situ hybridization, 100 um.

In *Vcar-eve* RNAi embryos that lost segments, *Vcar-en* expression localized in the stomodeum, the gnathal segments, and the telson (Figure 4Ci). In the case of embryos with partial segmental fusions, *en* was detected in the stomodeum, each partial segment, and the telson (Figure 4Cii). In knockdown germbands, we observed expression in the stomodeum, thoracic segments and telson, areas where *en* it is not normally expressed. Overall, although *Vcar-en* expression was clearly altered after *Vcar-eve* knockdown, we did not capture a stage at which PR-like loss of alternate *engrailed* stripes was observed.

### *Vanessa slp2* affects segmentation

We isolated two *Vanessa sloppy paired* genes that are orthologs of *Drosophila slp1* and *slp2*, each of which is expressed in a striped manner (Figure 3D-E). After *Vcar-slp1*-dsRNA1**-**injection, most embryos were wildtype-like (84%, n=475/564), some displayed delayed development (7%, n=39/564) or no development (9%, n=50/564). A second, non-overlapping dsRNA produced similar results (Table S1, Supplementary Fig S3B).

*Vcar-slp2* eRNAi produced a range of defects, with five distinct phenotypes comprising 64% (n=183/284) of dsRNA1-injected embryos and 35% (n=159/458) of dsRNA2-injected embryos (details in Table S1). The rest were wildtype-like, displayed delayed development, or failed to develop (comprising 36%, n=101/284 for dsRNA1 and 65%, n=299/458 for dsRNA2, Table S1). Three phenotypes were similar to those we observed for *Vcar-gsb* eRNAi: (1) head-like (Figure 4Di), (2) head and one set of prolegs (Figure 4Dii), and (3) torus (Figure 4Div). Other phenotypes include (4) head, legs and prolegs with the head capsule displaying wildtype-like stemmata, frons and antenna, one or two thoracic segments with leg-like structures, and five to six abdominal segments with prolegs (Figure 4Diii, arrowheads) and (5) a variety of partial fusions, with thoracic segments mainly affected. For example, we observed two fused thoracic segments with three legs and wildtype-like abdominal segments (Figure S2Bi asterisks indicating legs). Others displayed all thoracic segments, with two legs, and the first three abdominal segments fused (Figure S2Bii). Lastly, some displayed thoracic segments without legs, and some abdominal segments fused that produced loss of prolegs, often with only 2-3 prolegs present (Figure S2Biii).

Examination of *Vcar-en* expression in *Vcar-slp2*-dsRNA2-injected embryos revealed two main phenotypes: 1) very short germbands with defined head lobes but no segmental grooves and a V-shaped telson (Figure 4Ei); and 2) head lobes, fused gnathal, thoracic and first abdominal segments and a defined telson, with different degrees of fused segments in other parts of the embryo (Figure 4Eii-iii). *Vcar-en* expression was not uniform across these germbands, with some segments showing stripes (Figure 4Ei).

In *Drosophila*, *slp1* and *slp2* expression patterns are similar and have a redundant functional relationship in segmentation (Grossniklaus et al. 1992; Cadigan et al. 1994). We therefore tested if knockdown of both *Vcar-slp1* and *Vcar-slp2* would result in more severe segmentation phenotypes than either alone. Almost half (∼48%, n=209/433) of the embryos presented the five key *Vcar-slp2* phenotypes (Table S3). We did not observe more severe phenotypes for the mix of *Vcar-slp1* and *Vcar-slp2* dsRNAs, suggesting that *Vcar-slp2* is the main *slp* that functions in segmentation in the butterfly, but no PR-like function was seen.

### *Vanessa run* has a role in segmentation

To knock down *Vcar-run*, we designed two non-overlapping dsRNAs. dsRNA1 targets the region encoding the Runt domain; dsRNA2 targets the 3’ UTR (Fig S1G). *Vanessa* embryos injected with 1.5 uM of *Vcar-run*-dsRNA showed three main phenotypes (comprising 72%, n=446/617, of dsRNA1-injected embryos and 53%, n=333/624, of dsRNA2-injected embryos, Table S1). The remaining prehatchlings (comprising 28%, n=171/617, of dsRNA1-injected embryos and 47% n=291/624, of dsRNA2-injected embryos) appeared wildtype-like or displayed no development (Table S1). The most common segmentation phenotypes were: (1) Undefined mass of undeveloped tissue. In rare cases, it was possible to identify structures—like mandibles (Figure 4Fi, arrowhead) or a head capsule with stemmata, mandibles, and shortened body (Figure 4Fi)—within this mass. (2) Head that appeared wildtype-like, one thoracic segment with one set of legs, and 5-6 abdominal segments with 2-3 sets of prolegs (Figure 4Fiii-iv). The inconsistent presence of prolegs makes this phenotype difficult to classify as PR-like or not. (3) Partial segment fusions mainly in the thoracic region. In some cases, we observed two thoracic segments with legs and one without legs, and wildtype-like abdominal segments (Figure S2Ci). In others we observed fusions of thoracic segments and the presence of leg-like structures, and in some cases the fusions included the first abdominal segments with the presence of prolegs (Figure S2Cii-iii).

*Vcar-en in situ* hybridization in *Vcar-run-*dsRNA-injected embryos revealed low expression of *Vcar-en* (Figure 4Gi). In some cases, we observed diffuse *Vcar-en* expression in germbands with defined head structures and no segmental grooves (Figure 4Gii). In others, strong *Vcar-en* expression was observed in germbands with defined head structures, fused thoracic segments (Figure 4Giii-iv, arrowhead) and some defined abdominal segments with broad *Vcar-en* expression. Overall, most *Vcar-run* knockdown embryos displayed defects in thoracic and abdominal segment formation. Because it was difficult to determine the register of segmental deletions and fusions, we were unable to definitively determine whether *Vcar-run* has PR-like function but its striped expression pattern is consistent with a PR-role.

### An *Oncopeltus* PR- gene ortholog plays a novel role in *Vanessa*

In *Oncopeltus*, none of the *Drosophila* PR-gene orthologs play roles in PR-patterning (Reding et al. 2019) but *Oncopeltus* still follow a pair “rule” with genes *E75a* (Erezyilmaz et al. 2009) and *Blimp1* (Reding et al. 2024) having PR-expression and function. *Vcar-E75* expression was not detected in early embryos (Figure 5Ai-iii). Once the germband started to elongate, faint expression appeared at the stomodeum and midline (Figure 5Aiv-v). *Vcar-Blimp1* expression was detected as wide bands in the blastoderm, one in the anterior region and the second in the posterior region (Figure 5Bi, arrowheads), then in the germ primordium as two anterior bands while posterior band of expression remained (Figure 5Bii, arrowheads). Once the germband started to elongate, low expression in the protocephalon and three stripes in the posterior region were observed (Figure 5Biii, arrowheads). During germband elongation, two widely spaced stripes were observed anterior to the SAZ (Figure 5Biv, arrowheads). Later, *Vcar-Blimp1* expression was localized to the stomodeum and the midline region of posterior abdominal segments (Figure 5Bv, bracket).

**Figure 5.**
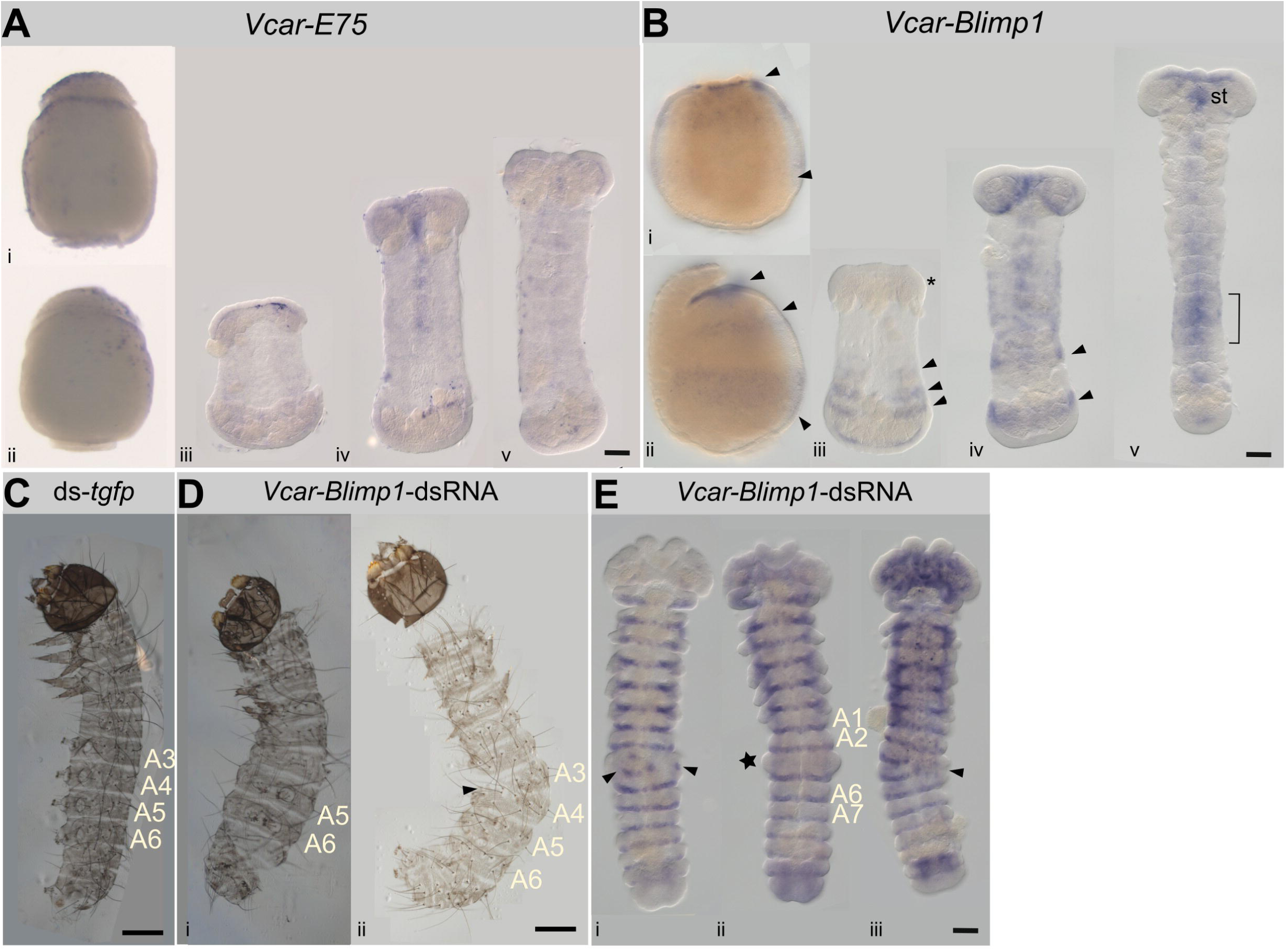
*Vcar-Blimp1* plays a novel role in segmentation. (**A**) *Vcar-E75:* expression not detected in (i) blastoderm or (ii) germ primordium. (iii-v) Expression marginally detectable during germband elongation. (**B**) *Vcar-Blimp1:* (i) one anterior and one posterior band in blastoderm; (ii) two anterior and one posterior band in germ primordium. (iii) Weak expression in head lobes and 3 stripes anterior of the SAZ. (iv) Midline expression, 2 widely spaced stripes (arrowheads). (v) Strong expression in abdominal segment primordia (bracket). (**C**) *Vanessa tgfp*-dsRNA-eRNAi, cuticular preparations were wildtype-like; segments A3-A6 indicated. (**D**) *Vcar-Blimp1*-eRNAi cuticular preparations: (i) loss of abdominal segments A3 and A4 or (ii) partial fusions of A3 and A4 (arrowhead). (**E**) *Vcar-en* expression in *Vcar-Blimp1*-eRNAi embryos. (i) Germbands with fused segments between segments A2 and A6; (ii) completely fused segments A3-A4 (star); (iii) partially fused A3-A4 segments. Images orientated anterior, top; posterior, bottom. Scale bars: cuticle preparations, 200 um; in situ hybridization, 100 um.

We performed *Vcar-E75* eRNAi experiments with two non-overlapping dsRNAs. *Vcar-E75* dsRNA1 and dsRNA2 match and extend slightly beyond the 5’ and 3’ portions, respectively, of the region encoding the Ligand Binding Domain (Figure S1K). After injection of *Vcar-E75*-dsRNA, most embryos appeared wildtype-like (dsRNA1, 93%, n=569/611, dsRNA2, 78%, n=244/312). A small number displayed delayed development or no development (see Table S1), similar to numbers seen for controls (Figure S3C). Thus, we found no evidence that *Vcar-E75* plays a role in segmentation.

*Vcar-Blimp1* knockdown was performed with two non-overlapping *Vcar-Blimp1*-dsRNAs, dsRNA1 matching the region encoding the SET domain and dsRNA2 targeting a portion of the region encoding the Zinc-Fingers (Figure S1L). Most embryos injected with *Vcar-Blimp1*-dsRNA showed complete or partial loss of segments A3 and A4, with no other segmental defects apparent (Figure 5Di; dsRNA1, 79%, n=431/545, dsRNA2, 56%, n=247/439, Table S1). In other cases, partial fusions of A3+A4 were observed on one side of the prehatchling but not the other, allowing us to count and definitively identify which segments were affected (Figure 5Dii, arrowhead). The remaining prehatchlings (dsRNA1, 21%, n=114/545, dsRNA2, 44%, n=192/439) appeared wildtype-like or displayed no development (Table S1).

*Vcar-Blimp1*-dsRNA-injected embryos displayed *Vcar-en* expression in every segment. In some cases, segmental fusions were observed (Figure 5Ei, arrowheads). Some germbands showed a bulging segment between segments A2 and A6 (Figure 5Eii, star). This enlarged segment appeared to fuse A3+A4+A5, based on germbands that showed fusion on only one side of the embryo but not the other (Fig. 5Eiii, arrowhead). These results suggest that *Vcar-Blimp1* functions in segmentation in a non-PR manner in *Vanessa*.

## Discussion

Here we described the early embryonic development of *Vanessa cardui* and expression patterns of *Vanessa gsb* and orthologs of the *Drosophila* and *Oncopeltus* PR-genes (Figs. 1, 2, 3, 5). All *Vanessa* PR-gene orthologs are expressed from blastoderm stage through germband elongation, except *Vcar-ftz-f1*. eRNAi knockdown suggests segment polarity-like function for *Vcar-gsb* and *Vcar-slp2*, classic PR-function for *Vcar-eve* while revealed *Vcar-run* knockdown embryos displayed fusion or loss of some or all segments, likely indicative of PR-function. Knockdown of *Vcar-Blimp1* produced fusions of abdominal segments A3 and A4 (Figs. 2, 4, S2, 5). Thus, some *Vanessa* segmentation genes share functions seen in *Drosophila* and other holometabolous insects, while others have different roles in embryonic development.

### *gsb* has segment polarity-like function in *Vanessa*

We showed previously that the *Pax3/7* family member *prd*, a PR-gene in *Drosophila*, is absent in Lepidoptera. However, we identified the presence of two genes from the *Pax3/7* family, *gsb* and *gooseberry neuro* (Gutiérrez Ramos and Pick 2025). *Bicyclus gsb* and *Vanessa gsb* are both expressed in late embryos in the mandibular segment (Matsuoka et al. 2023, Gutiérrez-Ramos and Pick 2025), similar to the localization of *prd* in the mandibular and maxillary segments in *Tribolium* (Coulcher and Telford 2012). *Bicyclus gsb* expression was also observed in the prolegs of late embryos, however we did not assess this stage in *Vanessa*. *Vcar-gsb* is expressed in segmental stripes prior to and during germband formation (Fig. 2), like segment polarity genes *wingless* (*wg*) and *en* in other lepidopterans (Kraft and Jackle 1994; Xu et al. 1997; Nakao 2010; Holzem et al. 2019; Hanly et al. 2021) and other arthropods examined (Patel et al. 1989; Nagy and Carroll 1994). In *Bombyx, wg* knockdown produced fused or truncated abdominal segments, in some cases the phenotype was similar to our *Vcar-gsb*-dsRNA torus phenotype (Yamaguchi et al. 2011; Zhang et al. 2015). In our functional experiments, *Vcar-gsb* knockdown phenotypes also produced severely truncated embryos, those composed primarily of head tissue and others with fused segments (Figure 2), similar to the head-only phenotypes seen for PR-gene knockdown in beetles (Choe et al. 2006; Xiang et al. 2017). Therefore, while we can conclude that *Vanessa gsb* plays a role in every segment, we are unable to rule out the possibility that it also has an earlier PR-like function.

### PR-gene expression patterns and function in other Lepidoptera

*Vcar-eve* and *Vcar-run* expression patterns are similar to those reported for their orthologs in *Bombyx. Bombyx-eve* presents and *Bombyx-run* are both expressed in PR-like stripes (Nakao 2010; Liu et al. 2008; Nakao 2015). *Bombyx-eve* eRNAi resulted in an asegmental phenotype (Xu et al. 1997; Nakao 2010), while we found classic PR-like loss of alternate body segments for *Vcar-eve* RNAi. *Bombyx-run* RNAi produced shortened abdomens (Liu et al. 2008, Nakao 2015) similar to *Vcar-run*. Thus, lepidopteran *eve* and *run* appear to share PR-roles with *Drosophila*. *Bombyx odd* showed an expression pattern similar to *Bombyx eve* and knockdown resulted in head-like phenotypes (Nakao 2015), with abnormal numbers of thoracic and abdominal segments, or loss of segments (Yamaguchi et al. 2011). While we did not test the function of *Vcar-odd*, its expression pattern is consistent with PR-like function.

The role of Lepidopteran *E75* orthologs during embryonic development has not been examined previously. Our results suggest that it does not play a role in segmentation in *Vanessa*. Lastly, *Bombyx Blimp1* expression and function were characterized in pupae and adults but no role in embryonic development had been reported (Wu et al. 2019). It will be interesting to see if the *Vcar-Blimp1* role in abdominal segmentation also applies to other butterflies and moths.

### Lepidopteran pair-rule gene orthologs within a broader context

Figure 6 summarizes gene expression and function in the dipteran *Drosophila*, three lepidopterans- *Bicyclus*, *Bombyx* and *Manduca*, two coleopterans— *Tribolium* and *Dermestes*, two hymenopterans—*Nasonia* and *Apis*, and the non-holometabolous hemipteran *Oncopeltus* for the 12 genes that have PR-function in at least one species, and two segment polarity genes, *wg* and *en*, that are expressed segmentally in all insect species examined (see references in Table S4).

**Figure 6.**
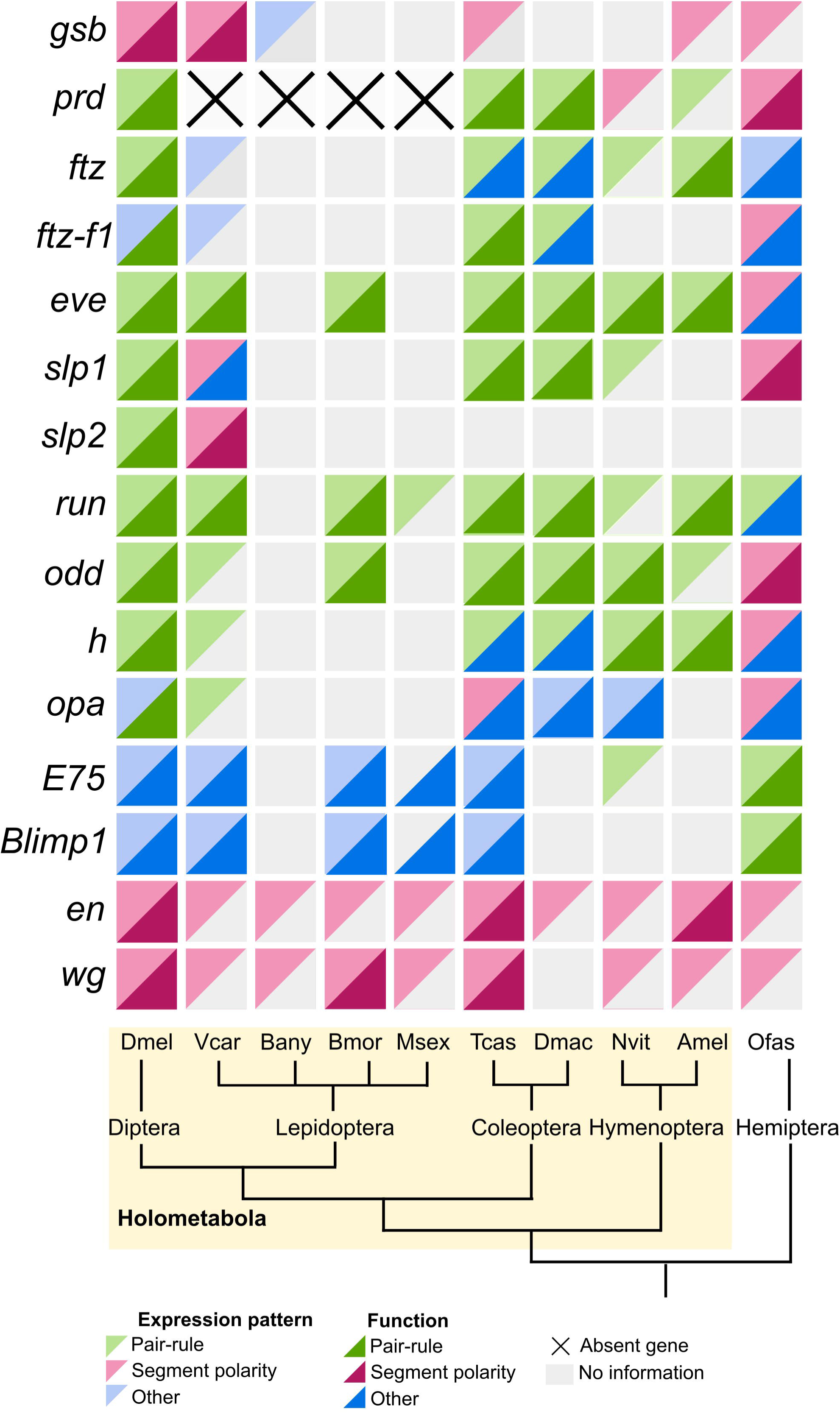
Variation in expression and function of insect segmentation genes. Summary of expression patterns and function of segmentation genes in selected species as indicated. Left, pair-rule and segment polarity genes (*wg* and *en*). Matrix, each row presents information about the expression pattern (light color) and the function of each gene (dark color) represented as triangles. Grey triangles if no information is available and X for absent genes. Each column represents insect species organized based on (bottom), phylogenetic relationships: Dmel – *Drosophila melanogaster*, Vcar – *Vanessa cardui*, Bany – *Bicyclus anynana*, Bmor – *Bombyx mori*, Msex – *Manduca sexta*, Tcas – *Tribolium castaneum*, Dmac – *Dermestes maculatus*, Nvit – *Nasonia vitripennis*, Amel – *Apis mellifera*, and Ofas – *Oncopeltus fasciatus*. Of the insect species used, all species are from the order Holometabola except *Oncopeltus fasciatus* (Hemiptera), an outgroup in which these genes have been studied. Summary of publications used to compile these data in Table S4.

In *Vanessa*, we showed that *gsb* is expressed segmentally and has segment polarity-like function, similar to *Drosophila* (Nusslein-Volhard and Wieschaus 1980), *Tribolium* (Aranda et al. 2008), *Apis* (Osborne and Dearden 2005) and *Oncopeltus* (Reding et al. 2019). We previously confirmed the loss of *prd* in Lepidoptera (Gutiérrez Ramos and Pick 2025). *prd* functions as a PR-gene in *Drosophila* (Nusslein-Volhard and Weisman 1980; Kilchherr et al. 1986) *Tribolium* (Maderspacher et al. 1998; Choe and Brown 2007), *Dermestes* (Xiang et al. 2015) and *Apis* (Osborne and Dearden 2005). However, in *Nasonia* and *Oncopeltus*, *prd* expression and function is segment polarity-like (Keller et al. 2010; Reding et al. 2019). *Vcar-ftz* is expressed as wide bands in the blastoderm, similar to *Tribolium* (Brown et al. 1994). *ftz* is expressed in a PR-manner in *Tribolium*, *Dermestes*, *Nasonia* and *Apis* (Wilson and Dearden 2012; Heffer et al. 2013; Xiang et al. 2017; Taylor and Dearden 2022) but is not required for PR-patterning in *Tribolium* or *Dermestes*. In *Vanessa* and all insects examined to date, *ftz* is expressed in the developing nervous system (Doe et al. 1988; Heffer et al. 2013; Reding et al. 2019). The expression of *Vcar-ftz-f1* in early embryos expression was not similar to other insects: in *Tribolium* and *Dermestes* the expression is PR (Heffer et al. 2013; Xiang et al. 2017) while it is segmental in *Oncopeltus* (Reding et al. 2019).

*eve* is the only gene with PR-expression and function across holometabolous insects examined to date, although more studies have examined *eve* than have examined several other PR-gene orthologs. In *Drosophila*, *Bombyx*, *Tribolium*, *Dermestes*, *Apis* and *Nasonia eve* is expressed in PR-stripes and knockdown resulted in the loss of alternate segments (Nusslein-Volhard and Wieschaus 1980; Macdonald et al. 1986; Patel et al. 1994; Binner and Sander 1997; Xu et al. 1997; Choe et al. 2006; Nakao 2010; Wilson et al. 2010; Wilson and Dearden 2012; Rosenberg et al. 2014; Xiang et al. 2017; Taylor and Dearden 2022). However, in *Oncopeltus*, like other *Drosophila* PR-orthologs, *eve* is expressed segmentally and functions segmentally (Liu and Kaufman 2005). Both *Vcar-slp1* and *Vcar-slp2* are expressed segmentally in the blastoderm, and *slp2* has segment polarity-like function. These observations contrast with the PR-expression of *slp* in *Drosophila*, *Tribolium*, *Dermestes* and *Nasonia*.

Similarly, PR-expression for *run* is likely highly conserved, having been documented for *Drosophila*, *Bombyx*, *Manduca*, *Tribolium*, *Dermestes*, *Nasonia*, *Apis*, and, to some extent in *Oncopeltus* (Gergen and Butler 1988; Kraft and Jackle 1994; Choe et al. 2006; Liu et al. 2008; Wilson and Dearden 2012; Rosenberg et al. 2014; Nakao 2015; Xiang et al. 2017; Taylor and Dearden 2022). In functional experiments we observed a likely PR-phenotype of loss of alternate segments, similar to *Drosophila* (Nusslein-Volhard and Wieschaus 1980), as well as a severe knockdown phenotype with only head-like structures observed in *Tribolium*, *Dermestes* and *Bombyx* (Choe et al. 2006; Liu et al. 2008; Xiang et al. 2017). Two other genes that have PR-expression in many holometabolous insects are *odd* and *h*. For *odd,* both expression and function in *Drosophila*, *Bombyx*, *Tribolium*, *Dermestes* and *Nasonia* have been reported as PR (Coulter et al. 1990; Choe et al. 2006; Yamaguchi et al. 2011; Rosenberg et al. 2014; Nakao 2015; Xiang et al. 2017; Taylor and Dearden 2022). However, changes have been observed for *h*, which has PR-function in *Drosophila, Nasonia* and *Apis* but knockdown of *Tribolium* or *Dermestes h* failed to reveal roles in segmentation (Ingham et al. 1985; Sommer and Tautz 1993; Aranda et al. 2008; Wilson and Dearden 2012; Rosenberg et al. 2014; Xiang et al. 2017; Taylor and Dearden 2022).

*Drosophila opa* functions as a PR-gene (Benedyk et al. 1994), however *opa* has shown variability in its early embryonic expression pattern and function among different insects. In *Vanessa*, we classified *opa* expression as PR-stripes, whereas in *Drosophila* it is expressed ubiquitously (Benedyk et al. 1994; Clark and Akam 2016). In *Tribolium* and *Oncopeltus, opa* is expressed segmentally; but no segmentation function was identified in *Oncopeltus* (Reding et al. 2019) and its function is unclear in *Tribolium* (Choe et al. 2017; Clark and Peel 2018). In *Dermestes* and *Nasonia, opa* expression and function are not PR (Xiang et al. 2017; Taylor and Dearden 2022).

*E75a* and *Blimp1* are PR-genes in *Oncopeltus* but little is known about their roles in embryonic development in holometabolous insects. It was observed that *Nasonia E75a* (Taylor and Dearden 2022) displays a PR-expression pattern, but we did not observe a clear pattern in the butterfly. While not PR, we observed a segmentation role for *Vcar-Blimp1* in abdominal segments A3 and A4. How *Blimp1* fits into a segmentation gene network in butterflies remains to be investigated.

In conclusion, the function and expression patterns of some genes important for segmentation in insects are conserved in a butterfly model system. *Vanessa gsb* appears to share with *Drosophila* a role as a segment polarity gene. While several of the orthologs of *Drosophila* PR-genes have PR-expression and/or function in *Vanessa,* others do not*. Vanessa* does not “make up for this deficit” by utilizing orthologs of the *Oncopeltus* PR-genes for PR-patterning. Interestingly, *Vcar-Blimp1* plays a different role in segmentation, being required for the formation of only two abdominal segments. Thus, this gene has changed function more than once during insect evolution, functioning as a PR-gene in *Oncopeltus* but playing no role in segmentation in *Drosophila*.

## Methods

### *Vanessa* rearing and embryo collection

The *Vanessa* colony is maintained in an incubator (Percival) at 25° C with a 12 hour light/dark cycle at 50% humidity. Larvae are maintained on an artificial diet (Multiple species diet, Southland Products) and adults are fed with a 50% diluted sports drink (Gatorade:tap water). Five-day old adults are exposed to a hollyhock plant for a defined period to collect embryos. Embryos are removed from the hollyhock leaves and grown in a petri dish until the desired time at 25° C.

### *Vanessa* gene ortholog identification and isolation

The annotated *Vanessa* genome annotation (PRJEB42869, ilVanCard2.1 (GCF_905220365.1)) was used to identify the *Drosophila* and *Oncopeltus* PR-gene ortholog sequences using tBLASTn. For gene isolation, primers (Table S5) were designed using the transcript sequences identified from NCBI. For RT-PCR, total RNA was extracted from embryos 16 to 24 hours AEL or from adult female posterior abdomens. For embryonic RNA, 100 uL of embryos were collected. Female posterior abdomens were used to extract adult female RNA. Total RNA was extracted using Trizol (Invitrogen) per manufacturer instructions and DNase-treated with TURBO DNA-free kit (Invitrogen). cDNA synthesis was done with NEB reagents using random primer mix. All PCR products were inserted into a pGEM vector (Promega) and Sanger-sequenced.

### Embryo fixation and nucleic acid staining

Embryos used for the nucleic acid staining were collected for 1-hour periods for the first 8 hours of development and an 8-hour collection was performed to examine embryos from 8 to 16 hours AEL. All embryos were allowed to develop in petri dishes under the same conditions as adults until the desired timepoint. For embryo fixation, a tube of ∼300 uL of embryos and 500 uL of water was boiled for 3 minutes, then moved to ice for 10 minutes. The chorion was manually removed in a Pyrex spot plate in tap water. The embryos were then fixed with 4% paraformaldehyde in phosphate-buffered saline with 0.1% Tween-20 (PBST) for 20 minutes by rocking, washed with PBST and dehydrated gradually into 100% methanol. Samples were stored at -20°C until use. For the nucleic acid stain, the embryos were rehydrated gradually into 100% PBST, then incubated at room temperature (RT) for 10 minutes with 50 ug/mL Hoechst-34580/PBST by rocking. They were rinsed three times with PBST and mounted on slides in 70% glycerol/PBST. Embryos were photographed with DIC and DAPI filter on a Zeiss Axio Imager M1 microscope and photos were arranged with ImageJ v.1.54f (Scheider et al. 2012) and Inkscape v.1.3.2.

### Embryo gene expression analysis

Embryo collections and fixation were done as above (as in **Embryo fixation and nucleic acids staining**) except that after boiling, embryos were washed with 1.5% bleach for 45 seconds, then were fixed by rocking in 4% paraformaldehyde in PBST for 20 minutes, washed with PBST and then dehydrated gradually into 100% methanol. Embryos were stored at −20°C until use. For in situ hybridization, embryos stored at -20°C were rehydrated into PBST and chorion and germbands were dissected from the yolk. The *in situ* hybridization protocol was done as described in Gutiérrez Ramos and Pick 2025. Embryo blastoderms were photographed with a Zeiss Discovery stereoscope in a Pyrex spot plate in PBST. Embryo germbands were mounted on a slide with 70% glycerol/PBST and photographed with DIC on a Zeiss Axio Imager M1 microscope; several photos were taken for each embryo that were then stitched together using Adobe Photoshop and arranged with ImageJ v.1.54f (Scheider et al. 2012) and Inkscape v.1.3.2.

### Embryonic RNA interference experiments

Synthesis of dsRNA and embryo microinjections 3 hours AEL were performed as described in Gutiérrez-Ramos and Pick 2025. For cuticle phenotype scoring, injected embryos were grown for 3 days, then boiled for 3 minutes, cooled and dissected from the chorion for examination. For imaging cuticles, each individual larva was mounted on a slide, covered with 100 uL of lactic acid, cover slipped, incubated overnight at 60°C, and photographed with DIC on a Zeiss Axio Imager M1 microscope; several photos taken for each larva were stitched together using Adobe Photoshop and arranged with ImageJ v.1.54f (Scheider et al. 2012) and Inkscape v.1.3.2. For gene expression pattern analysis, embryos were boiled, fixed and subjected to in situ hybridization 24 hours post injection, as described above.

## Data availability

The data underlining this article is available in supplementary material and GenBank PX977123-PX977135.

## Author contributions

Conceptualization: X.G.R. and L.P.; Methodology: X.G.R. and K.R.; Formal analysis: X.G.R.; Validation: X.G.R. and K.R.; Resources and supervision: L.P. Writing—original draft: X.G.R.; writing— review and editing: X.G.R., K.R. and L.P. Funding acquisition: L.P.

## Acknowledgments

We thank Luis Lorenzo Manzanares for suggestions for fluorescent microscopy photographs analysis. We thank Alys Cheatle Jarvela and Minh Le for comments on the manuscript.

