## Supplementary material for "Segmentation gene expression and function in *Vanessa cardui,* an emerging model for Lepidoptera": GutierrezRamos_etal_SuppFigs.docx

**Ximena Gutiérrez Ramos et al. Supplementary Figures and Figure Legends**

**Supplementary Figure S1. *Vanessa* pair-rule genes structures.** Schematic representation of the CDS of *Vanessa* PR-gene orthologs. Colored boxes denote protein domains and grey boxes under gene structures indicate the regions targeted by dsRNA. (**A**) *Vcar-gsb* isoform (x) 1 coding sequence (CDS) is 1416 base pairs (bp); Paired domain (PRD), Homeodomain (HOX) and an Octapeptide motif (OP) are shown. dsRNA1 is 402 bp; dsRNA2 is 305 bp. (**B**) *Vcar-ftz* complete cDNA is 2040 bp and its CDS is 1359 bp, encoding a complete HOX, as previously reported (Mulhair et al. 2023) and two motifs, LxxxL (LRAIL) and YPWM. dsRNA1 is 296 bp and dsRNA2 is 432 bp. (**C**) *Vcar-ftz-f1* complete cDNA is 8024 bp, with four predicted isoforms; their CDSs are *ftz-f1*-x1 with 1701 bp, *ftz-f1*-x2 with 1728 bp, *ftz-f1*-x3 with 1686 bp, and *ftz-f1*-x4 with 1626 bp. *Vcar-ftz-f1* encodes one zinc finger (ZF), a Ligand Binding Domain (indicated as NR-LBD) and the C-terminal AF-2. dsRNA1 is 785 bp and dsRNA2 is 463 bp. (**D**) *Vcar-eve* complete cDNA is 1045 bp. The 870 bp CDS encodes a HOX. dsRNA1 is 353 bp and dsRNA2 is 290 bp. (**E**) *Vcar-slp1* complete cDNA is 982 bp with a CDS 702 bp that encodes a Fork head (FH) domain. dsRNA1 is 355 bp and dsRNA2 is 378 bp. (**F**) *Vcar-slp2* complete cDNA is 1321 bp with a CDS 969 bp that encodes a FH domain. dsRNA1 is 399 bp and dsRNA2 is 406 bp. (**G**) *Vcar-run* complete cDNA is 1951; the 1230 bp CDS encodes a Runt (RUN) domain. dsRNA1 507 bp and dsRNA2 556 bp. (**H**) *Vcar-odd* complete cDNA is 1758 bp with a 1047 bp CDS that encodes four ZF domains located in the C-terminal half of the protein. (**I**) *Vcar-h* complete cDNA is 2035 bp; the CDS is 774 bp and encodes a basic helix–loop–helix (bHLH) domain, an Orange (O) domain and the WRPM motif in the carboxyl terminal. (**J**) *Vcar-opa* complete cDNA is 3560 bp and CDS is 1329 bp, that encodes four ZF domains. (**K**) *Vcar-E75* has four isoforms: *E75*-x1 CDS is 2854 bp, *E75*-x2 CDS is 2537 bp, *E75*-x3 CDS is 2723 bp, and *E75*-x4 CDS is 2553 bp. *Vcar-E75* encodes two nuclear receptor C4-type (NR C4-type) ZF domains and a nuclear receptor ligand-binding (NR-LBD) domains. dsRNA1 301 bp and dsRNA2 514 bp. (**L**) *Vcar-Blimp1* complete cDNA is 5923 bp and has three isoforms: *Blimp1*-x1 CDS is 3207 bp, *Blimp1*-x2 CDS is 3165 bp and *Blimp1*-x3 CDS is 3207 bp. *Vcar-Blimp1* encodes a Su(var)3-9, Enhancer-of-zeste and Trithorax (SET) domain and five ZF domains. dsRNA1 is 939 bp and dsRNA2 is 791 bp.

**Supplementary Figure S2. *Vcar-eve, Vcar-slp2* and *Vcar-run* partial fusion knockdown phenotypes.** Cuticle preparations of eRNAi embryos. (**A**) *Vcar-eve* knockdown phenotypes included head-like (i), and partial segmental fusions with the loss of two abdominal segments, presumably T1 and T3, leaving one thoracic segment with legs (dots) and the loss of one abdominal segment with one set of prolegs (arrowheads), presumably A4 (ii). Others present fusions of thoracic segments with the presence of three legs (dots) and wildtype-like abdominal segments (iii-iv). (**B**) For *Vcar-slp2* eRNAi knockdown, partial fusions mainly affected the thoracic segments. The most common phenotypes are: (i) two fused thoracic segments with three legs (stars) and wildtype-like abdominal segments, with the presence of 4 sets of prolegs (arrowheads); (ii) All thoracic segments, with just two legs (stars), and the first three abdominal segments fused; and (iii) thoracic segments present but lacking legs, and some abdominal segments fused that affected the presence of prolegs (arrowheads). (**C**) *Vcar-run* partial fusions mainly impacted thoracic segments. We observed the presence of two thoracic segments with their legs (stars) and one segment without legs with the rest of the abdominal segments wildtype-like arrowhead indicate prolegs (i). Others have thoracic segments fused with the presence of legs-like structures (stars) and fusions that cover the first abdominal segments; prolegs were present (arrowheads) (ii-iii). Images are orientated anterior, top; and posterior, bottom. Scale bar is 200 um.

**Supplementary Figure S3. Wildtype-like knockdown phenotypes for *Vcar-slp1* and *Vcar-E75*.** Cuticle preparations of eRNAi embryos. (**A**) *tgfp-*eRNAi hatchlings are wildtype-like; the head, three thoracic segments (T1-T3) and ten abdominal segments (A1-A10) were all observed. A3-A6 and A10 have prolegs. (**B-C**) Wildtype-like phenotypes were observed after eRNAi targeting (**B**) *Vcar-slp1* or (**C**) *Vcar-E75*. Arrowheads show the prolegs. Images are orientated anterior, top; and posterior, bottom. Scale bar is 200 um.

**Supplementary Figure S1**

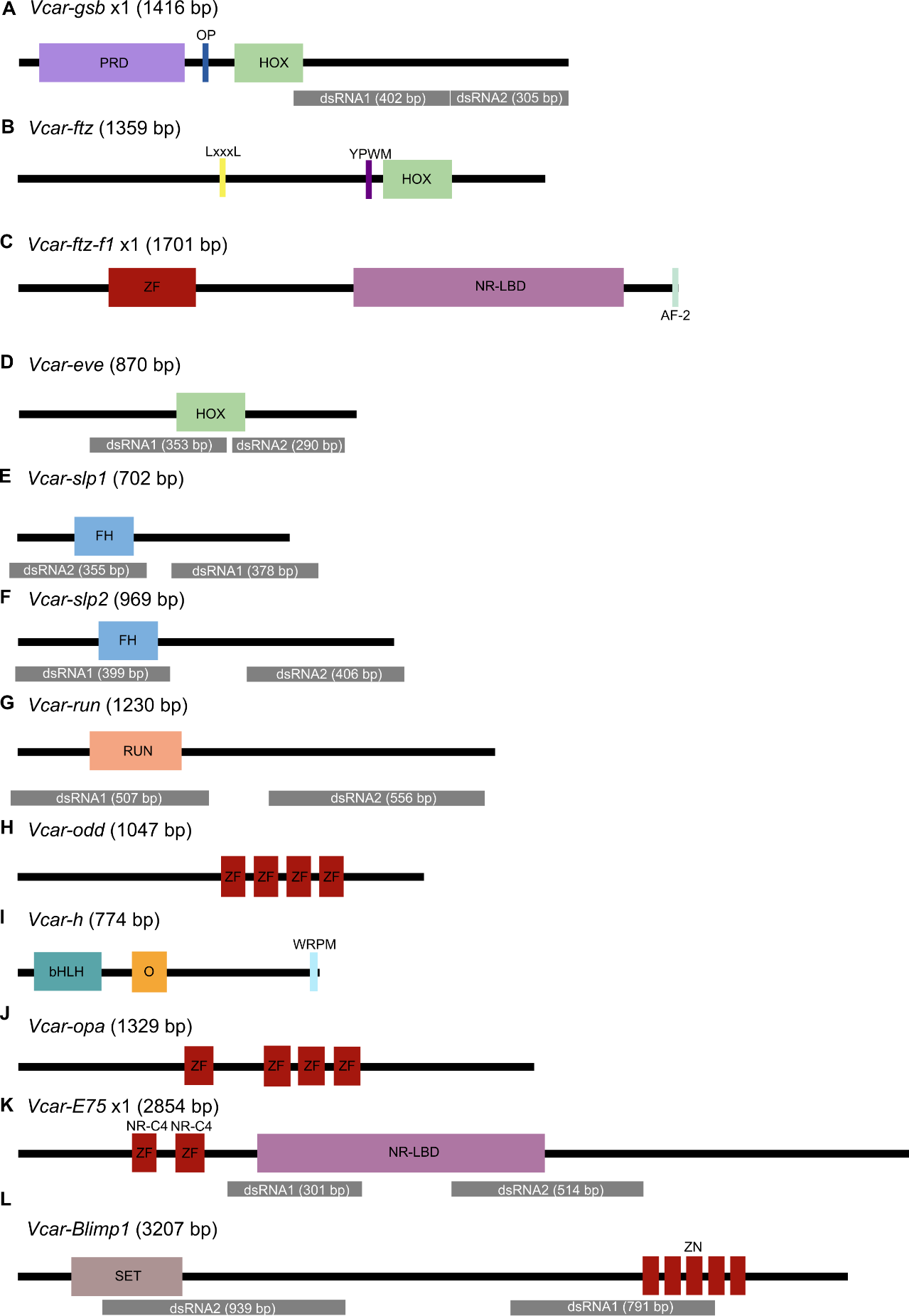

**Supplementary Figure S2**

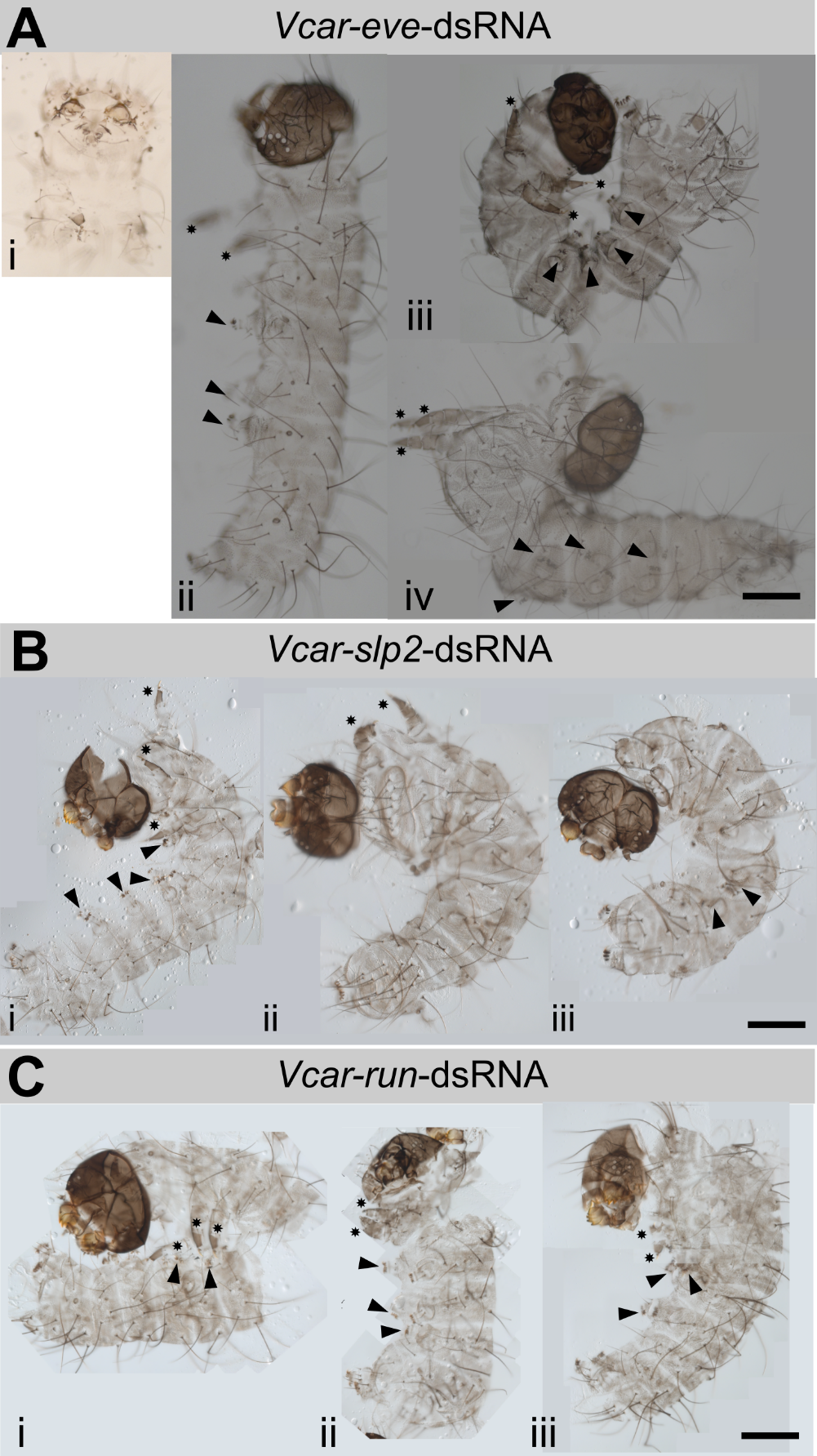

**Supplementary Figure S3**

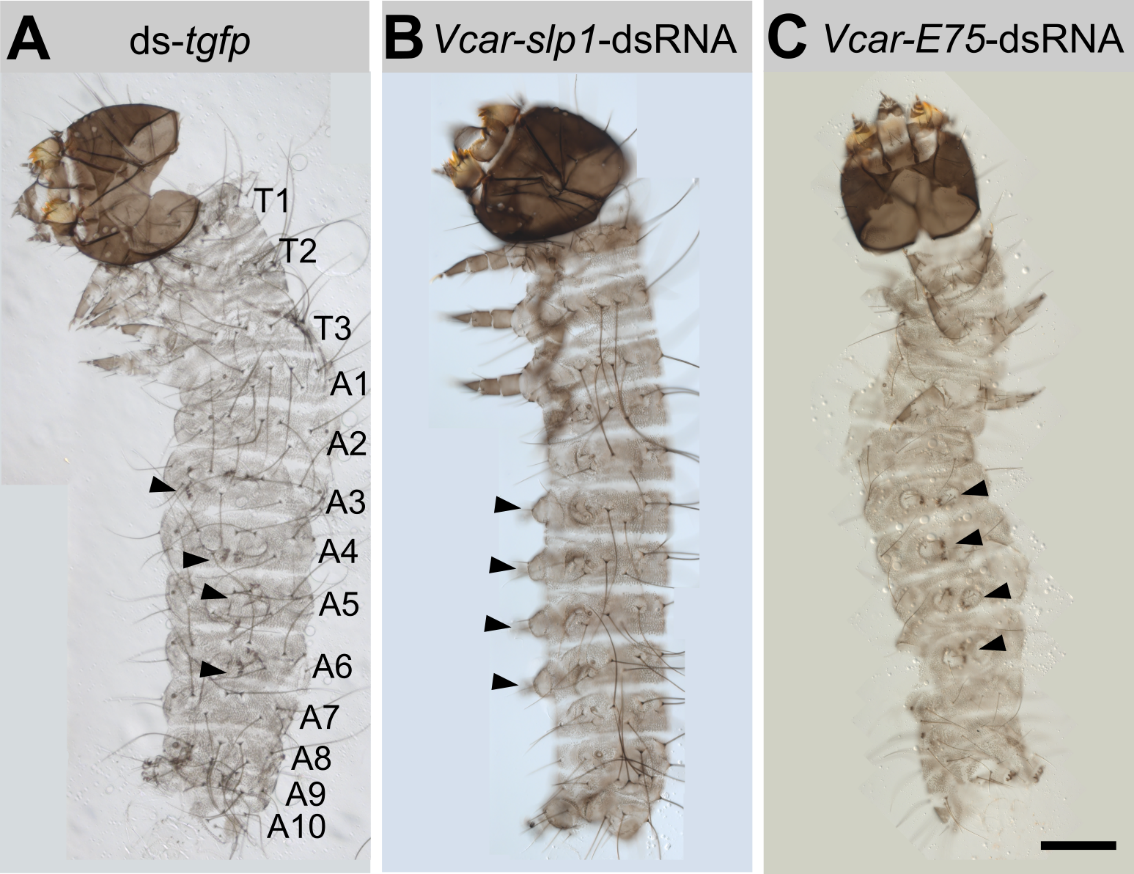
